# Loss of glia-neuronal interactions and age-dependent cell death in a *Drosophila* model of adult neurodegeneration

**DOI:** 10.1101/2024.05.20.595004

**Authors:** Unmila P. Jhuti, Edward M. Blumenthal

## Abstract

While glial dysfunction has been implicated in the development of multiple neurodegenerative diseases, the role of glial cell morphology in neurodegeneration is underexplored. In the fruit fly *Drosophila melanogaster*, mutants of the gene *drop-dead* (*drd*) exhibit adult neurodegeneration and extremely short lifespans. The morphology of one class of glia, the cortex glia (CG), is abnormal in *drd* mutants. In controls, the CGs form a continuous network that wraps around all neuronal cell bodies, but in *drd* mutants, individual CGs are stunted and the CG network is disrupted. These phenotypes are present on the first day of adulthood. Apoptosis is the central mechanism of cell death in *drd* mutants; widespread cell death is observed on the first day of adulthood and increases with age and is primarily neuronal. Apoptotic cells are found both within and outside of the remaining CG network, with significant variation in the distribution among individual brains. The degree of cell death and CG network breakdown in young adults could explain why *drd* mutant flies die within the first week of adulthood. The *Drosophila drd* mutant is a unique model of adult neurodegeneration that provides new insight into the breakdown in interaction between glia and neuronal cell bodies.

## Introduction

Glial cells are essential to the function of the nervous system, providing homeostatic support to neurons, maintaining immunological function, protecting brain tissues from injury and contributing to regeneration [1–5]. CNS-related neurodegenerative disorders were long thought to be solely neuron-dependent. However, recent research indicates that glial cell dysfunction is involved in several neurodegenerative diseases [6–10], suggesting that impairment of glial function can contribute to neuronal death and neurodegeneration (ND) [10–12].

The fruit fly *Drosophila melanogaster* is an excellent model organism for studying glial-related ND. *Drosophila* shares similar glial and neuronal functions with mammals [13,14]. The absence of different types of glial cells in flies causes severe brain defects and death [11,15–21]. A causative link between glial function and ND has already been demonstrated in *Drosophila* models of human disease. Glial dysfunction has been linked with major neurodegenerative diseases such as Alzheimer’s disease 1 and 2, SOD1-linked Amyotrophic lateral sclerosis 1, TDP-43 associated ALS10, Ataxia Telangiectasia and Huntington’s Disease [8–11,13,14,22,23].

The *Drosophila* central nervous system (CNS) structure is different than vertebrates in terms of the spatial localization of the cells. In the *Drosophila* CNS, neuronal cell bodies are in cortex, while neuronal processes and synapses lie within the neuropil regions [24,25]. The glia comprises 10-20% of the cells in the CNS and peripheral nervous system (PNS) [25,26]. There are predominately five different types of glial cells identified in the *Drosophila* CNS. They are cortex glia (CG), astrocyte-like glia (ALG), ensheathing glia, which can be subdivided into neuropil (NEG) and tract (TEG) ensheathing glia, and perineurial (PNG) and subperineurial glia (SPG) [25] (Fig. 1). The CGs reside in the cortex, and their processes form a honeycomb-like structure around the neuronal cell bodies. ALGs are sequestered in neuropil region with NEGs, while TEGs wrap the axons surrounding the neuropil. PNGs and SPGs are surface glia that form protective layers around the CNS and PNS [24,27–29].

**Figure 1:**
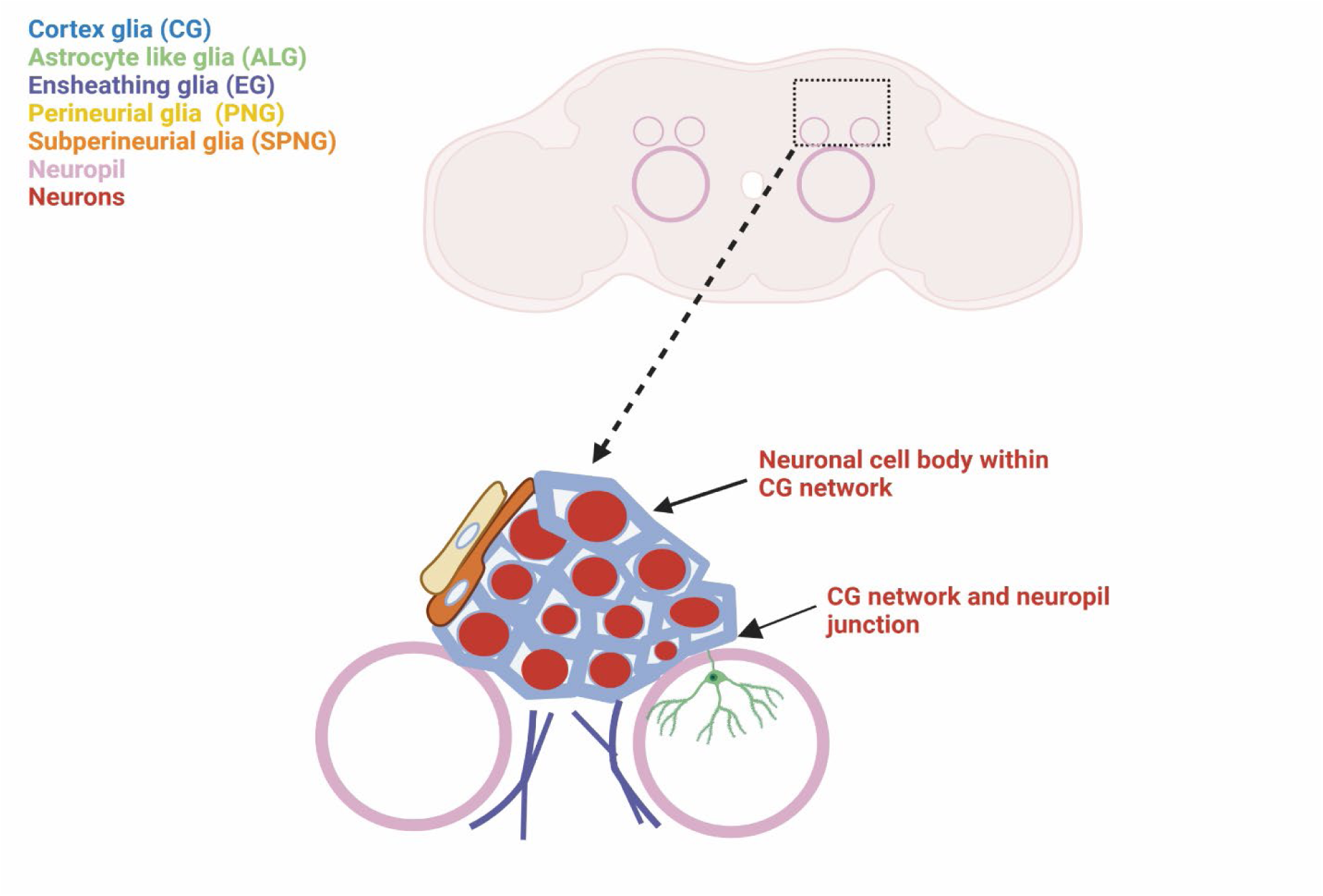
Schematic diagram of *Drosophila* glial subtypes showing their spatial distribution in the brain. CG network is continuous throughout the entire cortex region and does not extend into the neuropil.

In this study, we have studied the *Drosophila drop-dead* (*drd*) mutant, a unique model of ND. Flies mutant for *drd* exhibit ND and die as young adults within 4-8 days post-eclosion [30–33]. The *drd* gene is on the X chromosome and encodes an integral membrane protein predicted to have acyltransferase activity [33]. In addition to ND, mutation of *drd* disrupts eggshell integrity and digestive system structure and function [31,33–36]. Although the mutants show severe brain phenotypes with gross ND, the affected cell-type(s) and the mechanism remains unknown. The morphology of some glia has been reported to be altered in adult *drd* mutants, although the affected glial subtype(s) was not identified [31].

The current work describes a specific morphological defect in CGs in *drd* mutants. CG cells comprise around 20 to 35% of the total glial population in fly brain, with each CG wrapping processes around 40-100 neuronal cell bodies and the CGs together forming a continuous network throughout the cortex [24,29]. Insect CGs are structurally similar to vertebrate astrocytes and perform similar functions such as supplying nutrients to the neurons [25,37–40]. The CG network is formed starting in embryogenesis and develops its unique structure by the end of larval development [28,29]. Flies do not grow beyond larval L1 stage when their CGs are killed [41]. These observations suggest an essential role of CG in brain development or maintenance. We observe that in *drd* mutants, many of the CGs were smaller in size and lacked extensive processes. The absence of individual CG processes led to disruption of the CG network. Along with the CG phenotype, *drd* mutant brains showed age-dependent neuronal apoptosis, both within and outside the remaining CG network. This work establishes the *drd* mutant as a model for studying the link between the breakdown of glial association with neuronal cell bodies and ND in the adult brain.

## Results

### CGs in drd mutants show stunted morphology

The CGs in fly brain form a honeycomb-like structure that spans through the whole cortex region [24]. The MultiColor FlpOut (MCFO) technique, which stochastically labels a small number of cells in a given population, was used to visualize individual glia within the network in both controls and *drd* mutants [42]. In both genotypes, we observed CGs with similar size that wrapped processes around numerous neuronal cell bodies (Fig. 2a-2b). However, in mutant brains we observed smaller CGs as well (Fig. 2c-2d). These CGs lacked regular processes, were of smaller volume and did not wrap around the surrounding neurons (Fig. 2c-2d). We measured the volume of individual CGs, and as shown in Fig. 2e, the cell volumes in newly eclosed *drd* mutants fell into three populations, which we named regular, intermediate and stunted CGs (Fig. 2e). The mean volumes measured for regular CGs in controls and mutants were 3739.53 μm^3^ and 4195.51 μm^3^ respectively, whereas for stunted CGs, it was 204.98 um^3^ and for intermediate CGs, it was 718.45 μm^3^ (Fig. 2e). Overall, the CG volume in controls (mean= 3479.53 μm^3^) was significantly higher than in mutants (mean= 1629.71 μm^3^) (p=0.0002, Mann-Whitney test). The mean number of intermediate and stunted CGs labeled by MCFO in mutant brains was significantly higher than in controls; control brains typically had 6 regular, 0-1 intermediate and no stunted CGs, whereas mutant brains had 4 regular, 5-6 intermediate and 3-4 stunted CGs (Fig. 2f).

**Figure 2:**
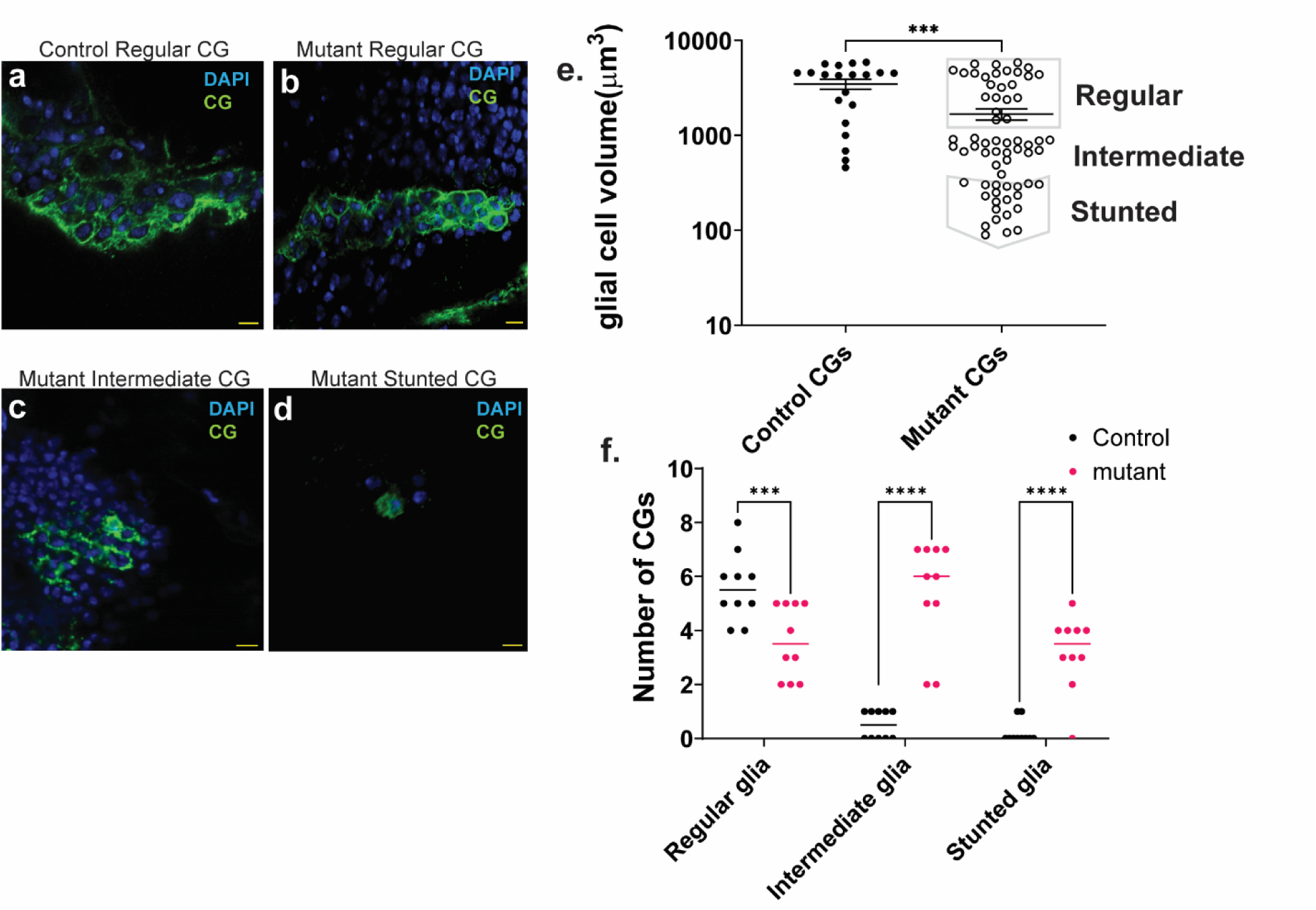
CGs show altered morphology in *drd* mutants. a. Single regular CG in a 0-day old adult control brain. b-d. Regular CG (b), intermediate CG (c) and stunted CG (d) in a 0-day old adult *drd* mutant brain. Scale bar: 10 μm. e. Comparison of individual CG volumes in mutant and control brains, showing the different populations of CGs, n= 20 cells from 10 control brains and 64 cells from 30 mutant brains, p= 0.0002 (Mann-Whitney test). f. Number of regular, intermediate and stunted CGs labeled by MCFO in each control and mutant brain. The number of regular CGs is significantly higher in controls, whereas the number of intermediate and stunted CGs are significantly greater in mutants, n= 10 control and 10 mutant brains, Fisher’s LSD test, p= 0.006 for regular, p<0.0001 for intermediate and stunted CGs.

### Other glial cell types in drd mutants show regular morphology

ALGs reside in the neuropil region and associate with synapses and processes [24]. The ALG processes extend from the cortex to the neuropil and form a contiguous network with other ALGs. Using MCFO, we generated control and *drd* mutant flies with individual labeled ALGs. We did not observe any differences in morphology between the two genotypes (Supplementary Fig. S1a-S1b). The mean volumes measured for control and mutant ALGs were 3410.68 μm^3^ and 3457.1 μm^3^ respectively (Supplementary Fig. S1c).

PNGs are distributed around the cortex regions in a tiled fashion [25]. Their shape and size can differ depending on their location in the brain. No difference in the morphology of individually labeled PNGs was observed between mutants and controls (Supplementary Fig. S2a-S2b). We have measured cell area instead of volume for PNGs and all other following glial cells because they are structurally flat compared to CGs and ALGs. The mean area for controls was 432.6 μm^2^ and for mutants was 419.5 μm^2^ (Supplementary Fig. S2c).

SPGs are very limited in number and they wrap the CNS with large processes in close proximity to the PNGs [43]. We did not observe any morphological differences between individually labeled SPGs in *drd* mutants compared with controls (Supplementary Fig. S3a-S3b). The mean area for controls was 577.7 μm^2^ and for mutants was 550 μm^2^ (Supplementary Fig. S3c).

Like ALGs, ensheathing glial cells are also found either in or adjacent to the neuropil. Although ensheathing glial cells in CNS and PNS are morphologically different [24], we focused on cells in CNS and looked at the morphology of two different types of ensheathing cells, TEGs and NEGs. No morphological differences between *drd* mutants and controls was observed for either cell type (Supplementary Fig. S4a-S4b; S5a-S5b). For controls, the mean area of TEGs was 1213.57 μm^2^ and for mutants, it was 1183.9 μm^2^ (Supplementary Fig. S4c). For controls, the mean area of NEGs was 978.3 μm^2^ and for mutants, it was 964.6 μm^2^ (Supplementary Fig. S5c).

### drd mutants show widespread age-dependent apoptosis

The cell death mechanism in *drd* mutants has not previously been identified. In order to identify the mechanism behind ND, we have performed acridine orange (AO) staining on 2-day old adult control and mutant brains and identified widespread presence of AO stained cells in mutant brains (2145 cells/brain) compared to control (41 cells/brain) (Fig. 3a-3c). Since AO staining is not specific for particular cell death mechanisms, we performed apoptosis-specific Death Caspase-1 (Dcp-1) staining in order to determine whether the cell death is due to apoptosis. We observed apoptotic events on the day of eclosion with increasing numbers as flies age. The mean number of Dcp-1 positive puncta in 0-day old and 2-day old mutants were 213 and 1031 respectively (Fig. 3d-3i, 3p-3q). Apoptosis was age-dependent as 2-day old brains showed significantly higher numbers of apoptotic puncta compared to 0-day old (Fig. 3r). The apoptotic puncta were widely distributed throughout the whole brain (Fig. 3d-3q).

**Figure 3:**
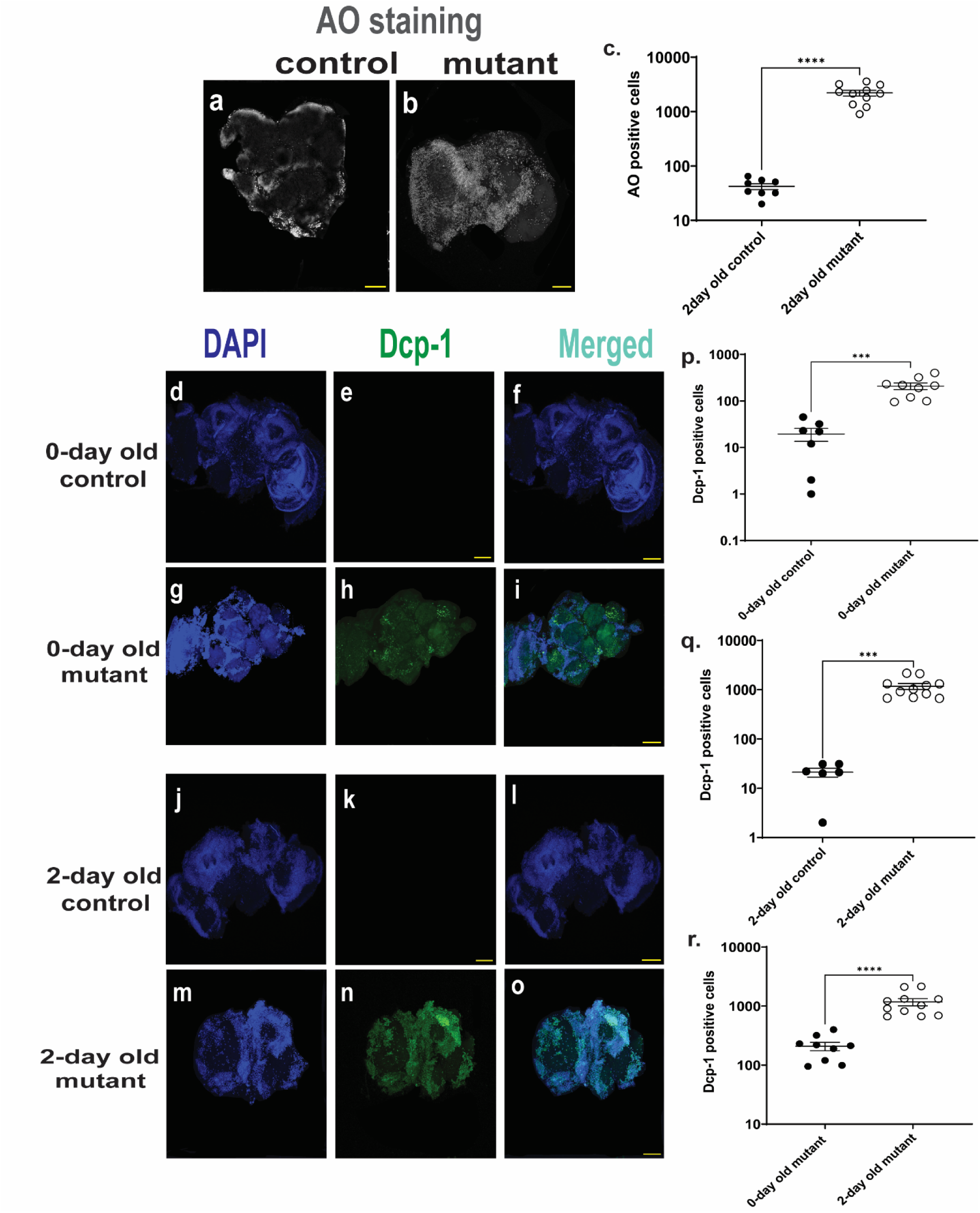
Apoptosis is observed in young and older *drd* mutants. Acridine orange (AO) staining showed widespread cell death in *drd* mutants. a. AO staining in 2-day old control brain. b. AO staining in 2-day old *drd* mutant brain. c. Comparison of the AO positive cells present in 2-day old mutant and control. n= 8 control and 11 mutant brains, p<0.0001 (Mann-Whitney test). d-f. DAPI and apoptotic Dcp-1 staining in 0-day old control brains. g-i. DAPI and apoptotic Dcp-1 staining in 0-day old mutant *drd* brains. j-l. DAPI and apoptotic Dcp-1 staining in 2-day old control brains. m-o. DAPI and apoptotic Dcp-1 staining in 2-day old mutant *drd* brains. Scale bar=100 µm. p-q. Comparison of the Dcp-1 puncta present in 0-day old and 2-day old mutant and control, p= 0.0002 in both cases (Mann-Whitney test). r. Comparison of Dcp-1 puncta in 0-day old and 2-day old mutant, p<0.0001 (Mann-Whitney test). n= 7 control and 9 mutant brains for 0-day old and n= 6 control and 12 mutant brains for 2-day old flies.

### Early adult apoptosis is primarily neuronal in drd mutants

From our previous experiment the cell death seems widely spread in mutant brains, but the identity of the dying cells in *drd* mutants was not determined. We performed double labelling in the brains of newly eclosed 0-day old flies with antibodies against neuron specific ELAV or glia specific REPO along with Dcp-1 to identify whether neurons or glia die first. We identified co-localization of Dcp-1 with neurons (Fig. 4a-4h in 0-day old mutants. Glial cells do not show colocalization with Dcp-1 in these brains (Supplementary Fig. S6a-S6h). Statistical analysis shows Dcp-1 puncta colocalization with ELAV in mutants is significantly higher compared to controls: mean Pearson’s rank coefficient for controls is 0.25, and for mutants is 0.72, where a value of 1 represents absolute colocalization and 0 represents no colocalization (Fig. 4i). The mean Dcp-1 colocalization with REPO was not significantly different in controls (0.20) compared with mutants (0.31) (Supplementary Fig. S6i).

**Figure 4:**
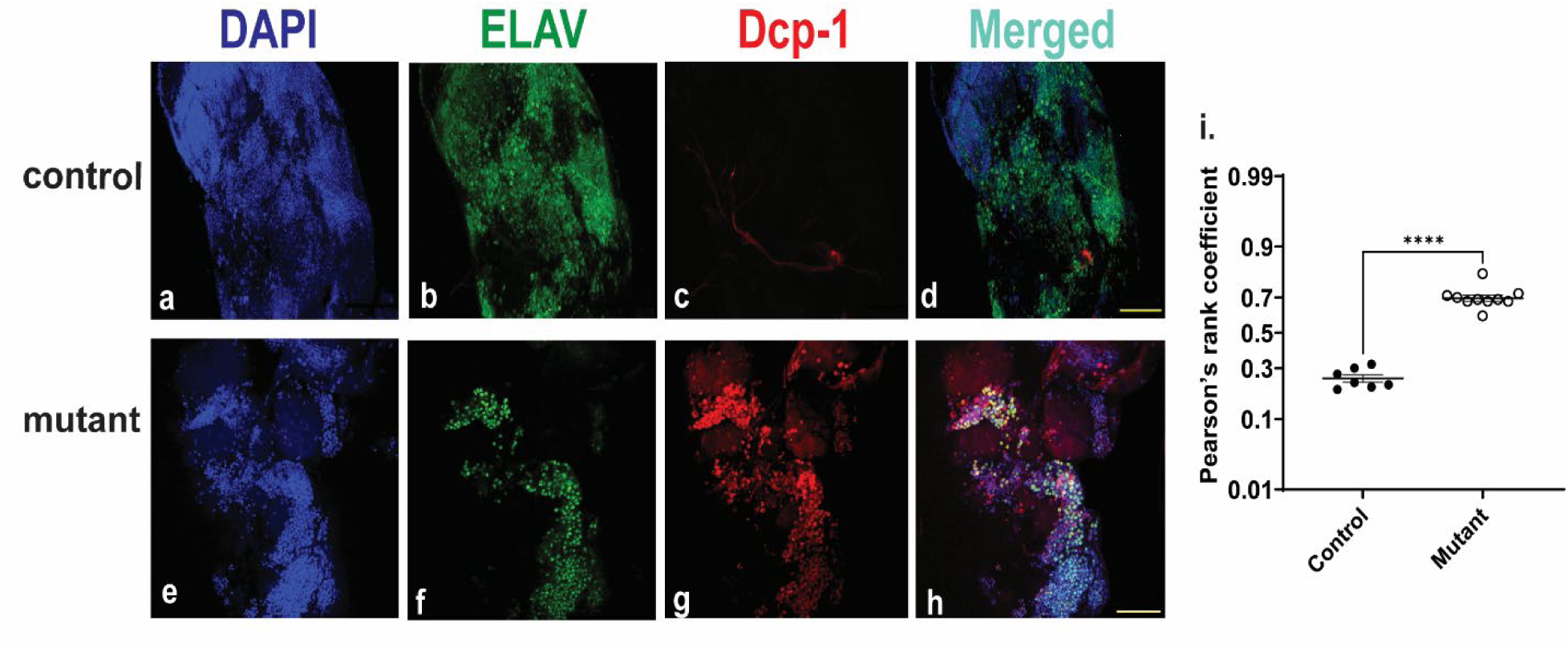
Neurons show apoptosis in 0-day old mutants. (a-h) co-localization of Dcp-1 and ELAV in brain cortex. Scale 100 µm. (i) Pearson’s rank coefficient for controls= 0.25, mutants= 0.72. n= 7 control and 10 mutant brains, p*<*0.0001 (Mann-Whitney test).

### drd mutants show discontinuous CG network and fewer viable cells

Since the CGs form a continuous network, having individual CGs with a stunted morphology may disrupt the integrity of the network. To visualize the complete network, we have driven GFP expression in all CGs. We observed a discontinuous CG network in mutant flies compared to controls (Fig. 5a-5h). The percentage of DAPI stained cells contained within the CG network was around 96% for controls and around 68% for mutants (Fig. 5i-5j). Overall, the mean number of DAPI stained cells was significantly lower in mutants (Fig. 5k). The reduction in the number of DAPI stained nuclei in mutant brains is consistent with the increased levels of apoptosis described above, and it raises the question of whether disruption of the CG network may lead to fewer number of viable cells in *drd* mutants.

**Figure 5:**
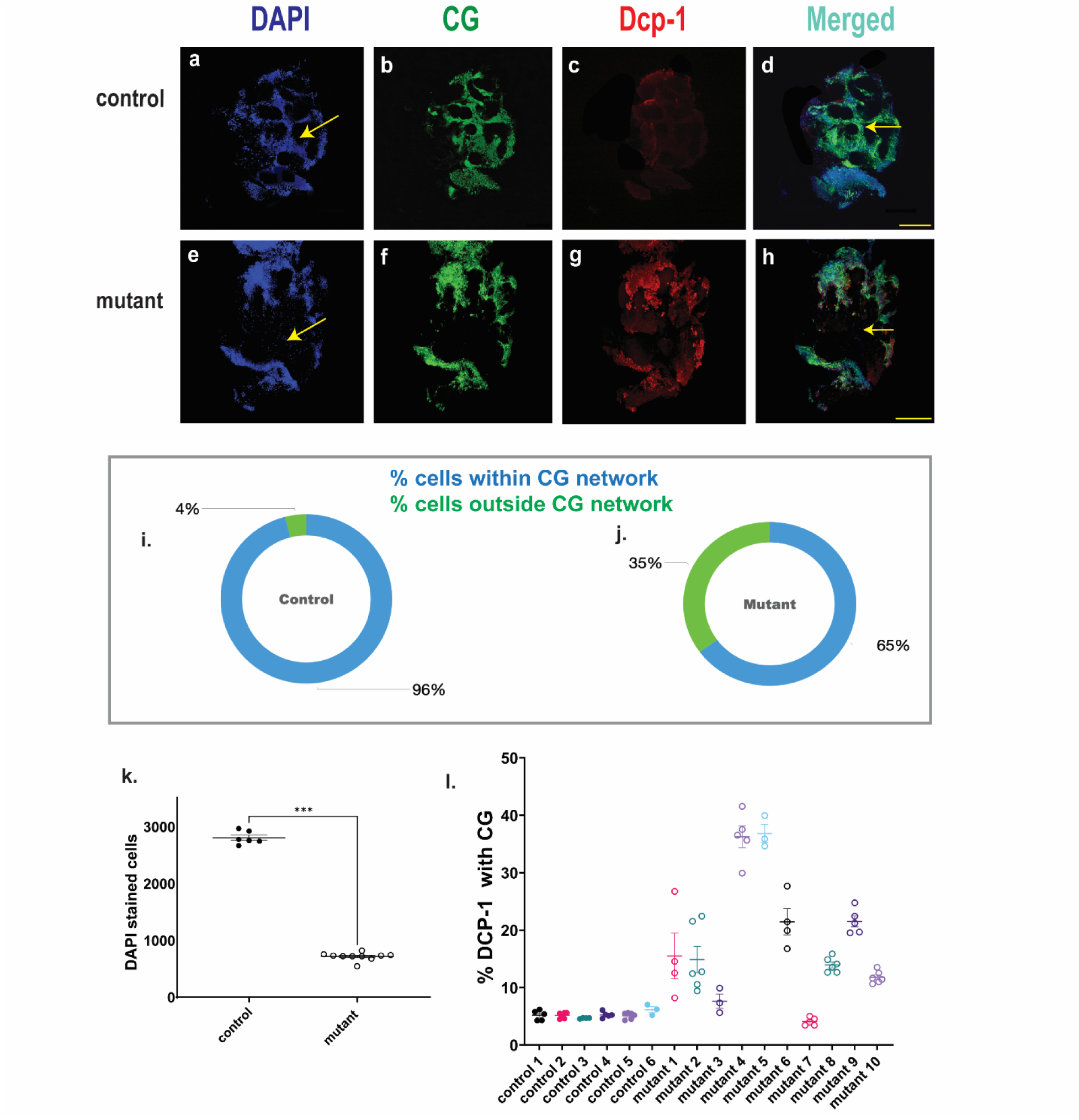
Fewer viable cells and a variable amount of apoptosis within a discontinuous CG network in 0-day old *drd* mutants. a-h. CG network (arrows indicate loss of CG network in mutant brain), DAPI and Dcp-1 in *drd* control and mutant. Scale 100 µm. i-j. Percentage of DAPI-stained nuclei colocalizing with the CG network is 96.43% in controls and 64.34% in *drd* mutants (Supplementary Table S1). k. The mean number of DAPI-stained nuclei is significantly higher in controls compared to mutants (arrows show loss of DAPI staining in mutant brain), mutants= 723 and controls= 2774, p*<*0.0001 (Mann-Whitney test). l. Percentage of Dcp-1 positive apoptotic cells within CG network. OBC analysis shows 5%-30% of Dcp-1 puncta within the CG network in mutants compared to <10% in controls. n= 6 controls and 10 mutant brains.

### drd mutants show increased apoptosis in broken CG network

We performed an object based colocalization (OBC) analysis to determine the fraction of Dcp-1 puncta localizing within the CG network (Dcp-1+CG). As shown in Fig. 5l, a variable percentage of apoptotic Dcp-1 was found within the CG network in *drd* mutant brains (5-30%), whereas in control brains, a consistently low percentage of apoptotic puncta were observed within the CG network. We performed a similar OBC analysis to quantify Dcp-1 staining outside of the CG network and observed no notable difference between control and mutant brains (Supplementary Fig. S7).

## Discussion

*drd* mutants exhibit apoptosis as the primary mechanism of cell death. Apoptosis is age-dependent, in that we observe significantly increased staining in 2-day old adults compared with newly eclosed adults. However, the extent of apoptosis in newly eclosed adults is so profound that the number of DAPI-stained nuclei is reduced by 75%. The cell death is not confined to a specific region of the brain, rather it is widespread both in young and old flies. We identified that the neurons, but not the glia, are the primary cells undergoing apoptosis in *drd* ND. This is the first detailed description of cell death and identification of a specific cell death mechanism in the *drd* mutant fly. *drd* mutant flies have a shorter median lifespan and show rapid aging [32,33,35,44] along with ND [30,31,35]. Neuronal apoptosis has been linked to nearly all reported neurodegenerative diseases, as well as to axon degeneration, CNS injury, and cellular stress [40,41,45–48]. However, the early, widespread, pervasive neuronal apoptosis observed in *drd* flies establish this mutant as a unique model of ND [30–33,35,44].

A second striking phenotype of adult *drd* mutant brains is the stunted morphology of many CGs. This phenotype is specific to CGs, as we did not observe any morphological changes in any other CNS glial class. Altered glial morphology is observed immediately after eclosion, which indicates the damage in CGs likely starts during metamorphosis. While the presence of morphologically stunted glia has previously been reported in *drd* mutants [31], the specific glial subtype affected was not identified. Along with stunted CGs, we observed a broken CG network in adult *drd* mutants, which could be explained by the accumulation of stunted CG processes and the failure of “regular” CGs to expand to maintain an intact network. Indeed, our observation that the percent of DAPI-stained nuclei lying within the CG network is reduced from 96% in control 0-day old adults to 64% in mutants suggests that at least one third of the CG network is eliminated in *drd* mutants by the start of adult life.

The existing literature on stunted CG morphology in adult flies is minimal. In larvae, a stunted CG morphology is observed upon disruption of vesicle fusion and recycling pathways, and this phenotype appears due to the lack of the neurotrophin Spätzle 3 [41]. Altered CGs and a compromised CG network were also observed in a ceramide phosphoethanolamine synthase *(cpes)*-null mutant model which eventually leads to photosensitive epilepsy [49]. CG and the overall glial network have a role in axon guidance as well. Disruption of CG network leads to the loss of axons in embryonic brains [50]. Disruption of the cortex and neuropil glia in larval brains altered the secondary axon growth and their trajectories [51]. Most models with compromised CGs either do not reach adulthood; or if they do, they show a significantly longer lifespan than *drd* mutants.

Adult *drd* mutants exhibit widespread neuronal apoptosis and a breakdown of the CG network. Given the association of stunted CGs and neuronal apoptosis in larvae [41], it can be suggested that the unwrapping of neurons caused by stunted glia potentially causes neuronal death. A simple model of this association would predict that apoptotic neurons would be exclusively localized outside of the remaining CG network. Our colocalization analysis showed that while most apoptotic puncta are found outside the CG network in both mutant and control brains, the percentage of apoptotic puncta within CG network is higher in mutants. This percentage also shows considerable variation among *drd* mutant brains—much higher variation than we observed in other metrics such as the number of stunted CGs or the number of DAPI-stained nuclei. Taken together, these data suggest a highly dynamic situation in the brains of young *drd* mutant adults in which the extent of CG network breakdown and neuronal apoptosis are changing rapidly. The spatial relationship between these two phenomena appears to be complex, possibly due to the short-term persistence of “unwrapped” neurons and the death of neurons within parts of the CG network that are structural intact but functionally impaired (Fig. 6).

**Figure 6:**
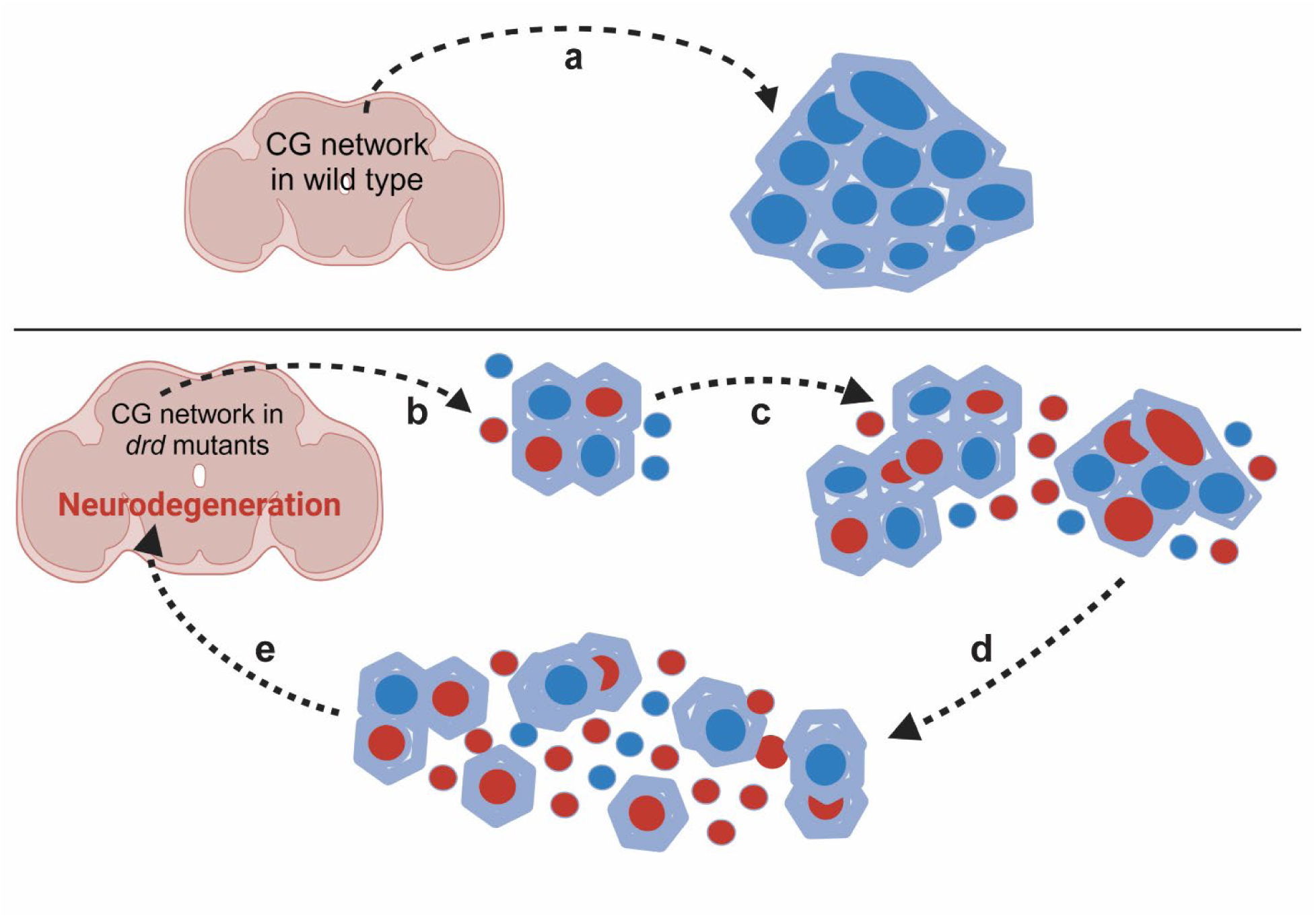
*Drosophila drd* model of ND. a. Wild-type brain displays regular CG morphology and healthy neurons (blue cells) within CG network. b. *drd* mutant brain displays stunted CG morphology and neuronal apoptosis (red cells). c. Neuronal apoptosis increases as stunted CGs are unable to wrap neurons. d. A large number of stunted CGs lead to a broken CG network with dying neurons within and outside the CG network. e. Continuation of age-dependent neuronal apoptosis and CG network breakdown subsequently leads to ND.

Stunted glial morphology [31] and rapid brain degeneration [32,35] have been the only reported hallmarks of ND in *drd* mutant brains. Our work showed the presence of stunted CGs, a broken CG network, and widespread neuronal apoptosis in newly eclosed mutant brains. Several simultaneous phenomena leading to ND suggest an essential role of *drd* in the fly brain. Interestingly, *drd* expression in neurons, glia or both cell types is not necessary for maintaining healthy brain function. Only tracheal expression of *drd* during metamorphosis is required for the flies to survive [35,44]. The reported requirement for drd expression during metamorphosis is consistent with our current finding that brain changes are well underway at the time of eclosion in *drd* mutants. Still unresolved is the connection between a loss of *drd* expression in respiratory tracheae and the breakdown of the CG network. This connection, however, is consistent with a known developmental interaction between tracheae and glia [52].

CGs provide neurons with structural support and perform several important functions such as phagocytosis, ion homeostasis and signaling [25,37–41,53]. These glia have an indispensable role because flies do not survive without CGs [40,41]. CGs and other glial cells also protect neurons from neuronal degeneration [45–47,54,55]. The *drd* mutant fly provides an important model for understanding the contribution of the CG network to neuronal survival and brain integrity. The extremely short lifespan and early widespread cell death accompanied by glial network degeneration make *drd* a unique model to study adult ND.

## Methods

### Fly stocks

Flies were maintained at 24°C on a 12h:12h light-dark cycle on standard cornmeal-molasses food except where indicated below. All fly stocks used in this study are listed in the Supplementary Methods. For acridine orange, Dcp-1, colocalization (Dcp-1+ELAV and Dcp-1+REPO) experiments, *w drd^lwf^/FM7a* female flies were used as controls and *w drd^lwf^/ w drd^lwf^* females were used as experimental flies.

### MCFO labeling of individual glia

Protocol was adapted from [56]. We developed *w drd^lwf^, hs-FLPG5 PEST/FM7i; UAS-HA-UAS-V5-THS-UAS-FLAG* by standard crossing and recombination from the following stocks- *w drd^lwf^ /FM7a, FM7i / X^X*, BDSC 62118 and BDSC 64093. After generating the *drd* MCFO flies, we crossed the stock with each glial-specific Gal4 line listed in the stock table (supplementary methods). Male offspring from the crosses were collected as experimental flies. For generating controls, the glial Gal4 lines were crossed with BDSC 64085 and male progeny were collected. All MCFO crosses were kept at 18°C. P12 pupal progeny from the crosses were collected, heat shocked for 30 s at 37°C and maintained at 24°C following heat shock. At 0 and 2 days post-eclosion, brains were dissected in Schneider’s medium and fixed with 4% paraformaldehyde in PBS. After fixation, brains were washed 3x for 15 min in adult brain wash solution (0.5% bovine serum albumin, 1% Triton X-100 in PBS) and blocked for 1 hr in blocking solution (3% normal goat serum, 3% normal donkey serum, and 1% Triton X-100 in PBS). Brains were incubated overnight at 4°C with rat anti-FLAG (DYKDDDK) primary antibody (Agilent technologies) diluted in adult brain wash solution (1:100). The following day, tissues were washed 3x for 1 hr in adult brain wash solution and incubated overnight at 4°C or for 4 hr at room temperature with AlexaFluor488 anti-rat (1:250) (Invitrogen) and DAPI (1:250) (Biotium) diluted in wash solution. Brains were washed 3x for 1 hr in adult brain wash solution, followed by a final wash in PBS overnight at 4 °C or for 1-2 h at RT. All the above steps were performed at room temperature except where noted. Brains were mounted in Vectashield (Vector Laboratories) on glass slides with spacers.

### Acridine orange staining

Brains were dissected from 2-day old adults in PBS and incubated in 1.6 × 10^-6^ M acridine orange (ThermoFisher) for 5 min at room temperature. Brains were washed quickly 3-4x in PBS, mounted in Vectashield, and imaged.

### Dcp-1 immunostaining

The immunostaining protocol was similar to that used for MCFO above. Primary antibody used was cleaved Drosophila Dcp-1 rabbit antibody (1:100) (Cell Signaling Technologies) and secondary antibody was AlexaFluor488 goat anti-rabbit (1:250) (Invitrogen) with DAPI (1:250) (Biotium). To observe the relationship between apoptosis and the CG network, the wrapper>myrTomato stock was crossed with *w drd^lwf^ /FM7a* or w* females to generate mutant and control males respectively. MyrTomato staining was observed directly at 546nm in fixed samples following anti-Dcp-1 immunostaining.

To observe colocalization of Dcp-1 with neurons and glia, brains of 0-day old adult flies were double immunostained against Dcp-1 (1:250) and either ELAV (mouse Elav-9F8A9 antibody, 1:100, Developmental Studies Hybridoma Bank (DSHB)) or REPO (mouse 8D12 anti-Repo antibody, 1:100, DSHB). Secondary antibodies used were AlexaFluor 488 goat anti-mouse (1:250, Invitrogen) and AlexaFluor 568 goat anti-rabbit (1:250, Invitrogen) with DAPI (1:500).

### Imaging

For confocal microscopy, a Nikon A1R system and NIS Elements AR 5.21.02 imaging software was used. Objectives used were: 20X, 60X and 100X with numerical aperture of 0.75. Optical sections were taken in intervals of 0.50 or 0.75μm.

### ImageJ analysis

Cell counts, volumes and areas were analyzed using Fiji ImageJ version 2.14.0/1.54f [57]. For volume analysis, stack slices were segmented and deconvoluted. Macro code used for volume analysis is shown in Supplementary Text S1. For area measurement, images were subjected to threshold and a region of interest (ROI) was manually selected. The data was then analyzed using Analyze-> measure. For counting cells, both stacks and maximum projections were used. ROI was manually selected when necessary. The images were deconvoluted, background was subtracted and threshold was set. The data was analyzed using Analyze-> Analyze Particles. For area measurement, images were subjected to threshold and a region of interest (ROI) was manually selected. The data was then analyzed using Analyze-> measure. For counting cells, both stacks and maximum projections were used. ROI was manually selected when necessary. The images were deconvoluted, background was subtracted and threshold was set. The data was analyzed using Analyze-> Analyze Particles.

To measure colocalization, the PTBIOP plugin was used with BIOP JACoP option was selected. Individual stack was split and the channels of interest (green and red) were subjected to analysis. Threshold was set and Pearson’s correlation was calculated. For object based colocalization, stacks were selected and channels of interest (green and red or green and blue) were split for analysis. Threshold was set and total area of Dcp-1, DAPI and CG were measured. AND and SUBTRACT functions from image calculator were selected in order to calculate area of Dcp-1 AND CG, Dcp-1 SUBTRACT CG, DAPI AND CG. Area of the AND and SUBTRACT data were divided by total Dcp-1 or total DAPI data in order to get a percent area.

### Statistics and data analysis

Data were graphed and analyzed using GraphPad Prism 9.5.1 [58]. 2-way ANOVA and Tukey’s multiple comparison test were performed to compare the stunted, intermediate and regular CG numbers in MCFO brains. Mann-Whitney non-parametric tests was used to compare cell volumes and areas, Dcp-1, AO, colocalization datasets. The object based colocalization (AND and SUBTRACT) was performed to identify Dcp-1 colocalization within CG network.

## Acknowledgements

Stocks obtained from the Bloomington Drosophila Stock Center (NIH P40OD018537) were used in this study. The anti-ELAV and anti-REPO monoclonal antibodies, developed by Drs. Gerald Rubin and Corey Goodman, respectively, were obtained from the Developmental Studies Hybridoma Bank, created by the NICHD of the NIH and maintained at The University of Iowa, Department of Biology, Iowa City, IA 52242. Support was provided by Marquette University and the Northwestern Mutual Data Science Institute. BioRender.com was used to create Figure 1 and Figure 6. We thank Dr. Jaeda Coutinho-Budd for Drosophila stocks and Drs. Lisa Petrella and Anita Manogaran for helpful discussions.

## Author contributions

UPJ and EMB conceptualized and designed the project and experiments. UPJ conducted the experiments, analyzed data, created figures and drafted the manuscript. UPJ and EMB revised the figures and manuscript.

## Data availability statement

The datasets generated during the current study are available in the e-Publications@Marquette repository, https://epublications.marquette.edu/blumenthal_lab/. Fly stocks are available upon request.

