## Supplemental material for "Loss of glia-neuronal interactions and age-dependent cell death in a *Drosophila* model of adult neurodegeneration"

**Supplementary methods**

*Table of fly stocks used in this study*

| Stock | Source | Purpose |
|---|---|---|
| *w** | Lab stock | Control stock |
| *FM7i, P{w[+mC]=ActGFP}JMR3/C(1)DX, y[1] f[1]* | Bloomington *Drosophila* Stock Center (BDSC) stock 4559 (FBst0004559) | FM7i-GFP / X^X, for crossing schemes |
| *FM7i /C(1)DX, y[1] f[1]* | BDSC stock 5263 (FBst0005263) | FM7i / X^X, for crossing schemes |
| *w drd$^{lwf}$ /FM7a* | Lab stock | *drd* mutant stock |
| *w[1118]; PBac{y[+mDint2] w[+mC]=10xUAS(FRT.stop)myr::smGdP-HA}VK00005 P{y[+t7.7] w[+mC]=10xUAS(FRT.stop)myr::smGdP-V5-THS-10xUAS(FRT.stop)myr::smGdP-FLAG}su(Hw)attP1* | BDSC stock 64093 (FBst0064093) | For creation of *drd* mutant MCFO stock |
| *w[1118] P{y[+t7.7] w[+mC]=hs-FLPG5.PEST}attP3; PBac{y[+mDint2] w[+mC]=10xUAS(FRT.stop)myr::smGdP-HA}VK00005 P{y[+t7.7] w[+mC]=10xUAS(FRT.stop)myr::smGdP-* | BDSC stock 64085 (FBst0064085) | control MCFO parental stock |

| | | |
|---|---|---|
| *V5-THS-10xUAS(FRT.stop)myr::smGdP-FLAG}su(Hw)attP1* | | |
| *w[1118] P{y[+t7.7] w[+mC]=hs-FLPG5.PEST}attP3* | BDSC stock 62118 (FBst0062118) | For creation of *drd* mutant MCFO stock |
| *w[1118]; P{y[+t7.7] w[+mC]=GMR54H02-GAL4}attP2* | BDSC stock 45784 (FBst0045784) | wrapper-Gal4 |
| *w[1118]; P{y[+t7.7] w[+mC]=GMR86E01-GAL4}attP2* | BDSC stock 45914 (FBst0045914) | ALG-Gal4 |
| *w[1118]; P{y[+t7.7] w[+mC]=GMR85G01-GAL4}attP2* | BDSC stock 40436 (FBst0040436) | PNG-Gal4 |
| *w[1118]; P{y[+t7.7] w[+mC]=GMR54C07-GAL4}attP2* | BDSC stock 50472 (FBst0050472) | SPG-Gal4 |
| *w[1118]; P{y[+t7.7] w[+mC]=GMR75H03-GAL4}attP2* | BDSC stock 39908 (FBst0039908) | NEG-Gal4 |
| *w[1118]; P{y[+t7.7] w[+mC]=GMR56F03-GAL4}attP2* | BDSC stock 39157 (FBst0039157) | TEG-Gal4 |
| *Sp/CyO; wrapper>QF2>Gal4, QUAS-myrTomato 3X HA, UAS-CD8:GFP/TM3[Sb, Tb-RFP]* | Jaeda Coutinho-Budd | wrapper>myrTomato (This stock was used in the absence of FLP to label the entire CG network with Tomato) |

**Supplementary Figure S1**

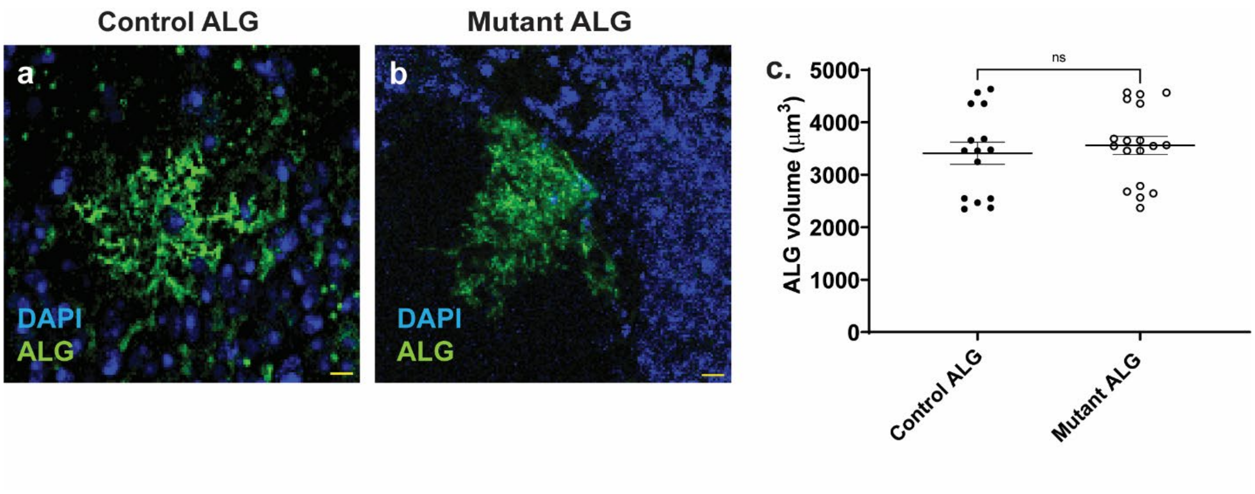


**Supplementary Figure S1:** ALGs show regular morphology in *drd* mutants. a. A single ALG in a 2-day old control brain. b. A single ALG in a 2-day old adult *drd* mutant brain. Nuclei were stained with DAPI (blue), and anti-FLAG was used to label ALGs (green). Scale bar=10 μm. c. Comparison of ALG cell volumes in mutant and control. n= 15 cells from 12 control brains and 18 cells from 12 mutant brains, Mann-Whitney test, p=0.46.

**Supplementary Figure S2**

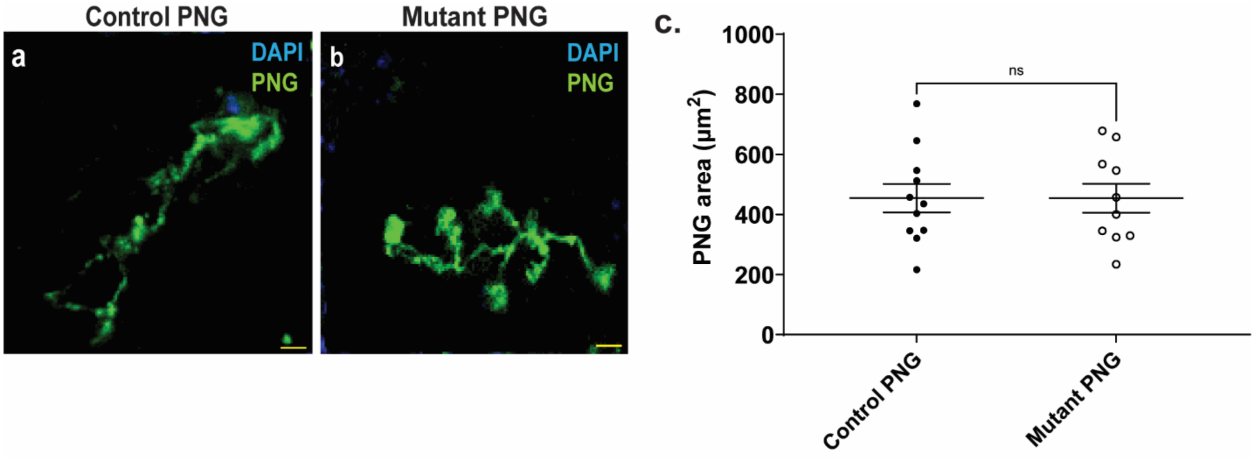


**Supplementary Figure S2:** PNGs show regular morphology in *drd* mutants. a. A single PNG in a 2-day old control brain. b. A single PNG in a 2-day old adult *drd* mutant brain. Nuclei were stained with DAPI (blue) and anti-FLAG was used to label PNGs (green). Scale bar=10 μm. c. Comparison of PNG cell areas in mutant and control. n= 11 cells from 7 control brains and 10 cells from 7 mutant brains, Mann-Whitney test, p=0.9.

**Supplementary Figure S3**

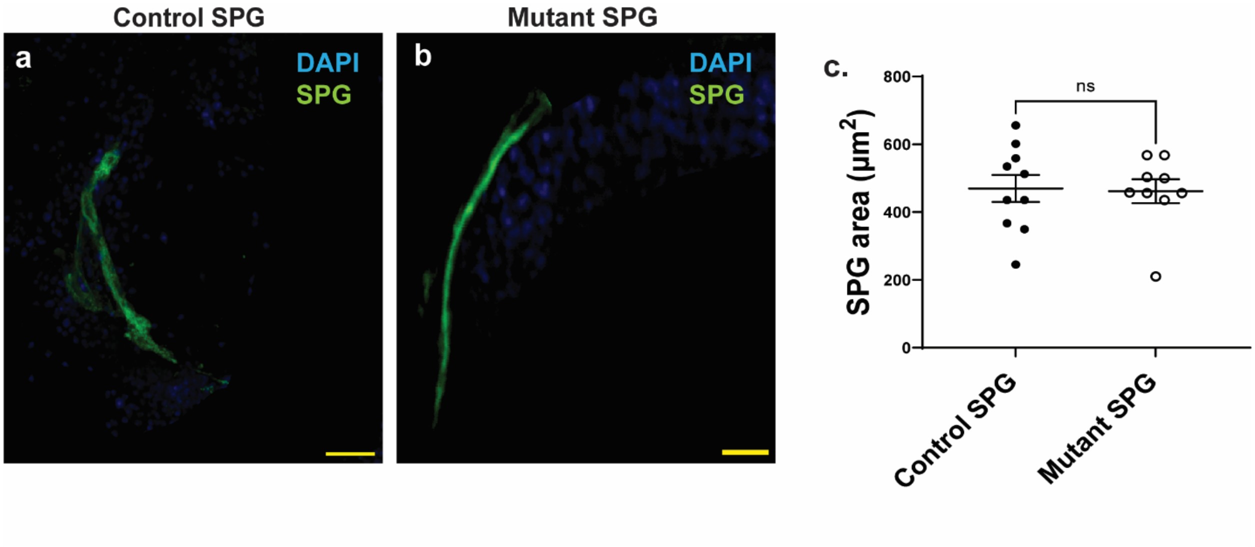


**Supplementary Figure S3:** SPGs show regular morphology in *drd* mutants. a. A single SPG in a 2-day old adult control brain. b. A single SPG in a 2-day old adult *drd* mutant brain. Nuclei were stained with DAPI (blue) and anti-FLAG was used to label SPGs (green). Scale bar=10 μm. c. Comparison of SPG cell areas in mutant and control. n= 10 cells from 10 control brains and 9 cells from 7 mutant brains, Mann-Whitney test, p=0.89.

**Supplementary Figure S4**

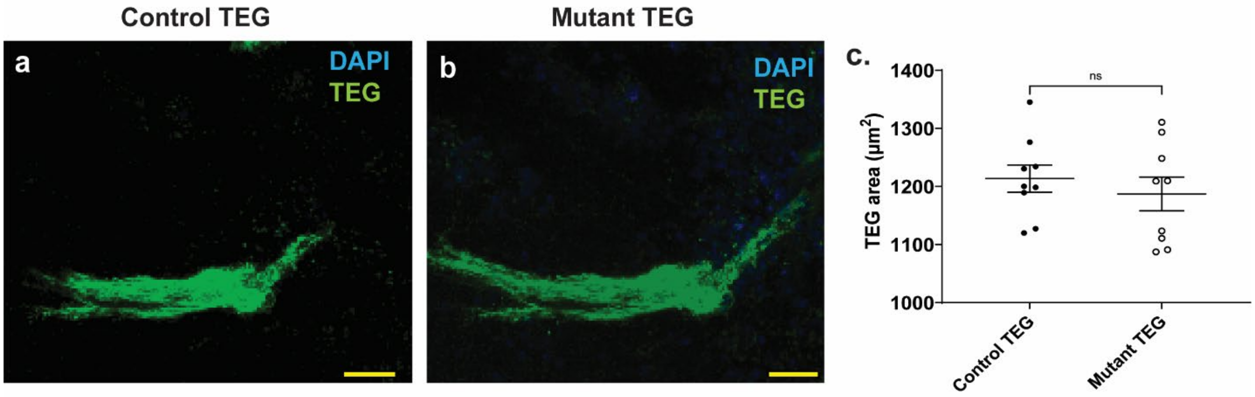


**Supplementary Figure S4:** TEGs show regular morphology in *drd* mutants. a. A single TEG in a 2-day old adult control brain. b. A single TEG in a 2-day old adult *drd* mutant brain. Nuclei were stained with DAPI (blue) and anti-FLAG was used to label TEGs (green). Scale bar=10 μm. c. Comparison of TEG cell areas in mutant and control. n= 9 cells from 11 control brains and 9 cells from 11 mutant brains, Mann-Whitney test, p=0.8.

**Supplementary Figure S5**

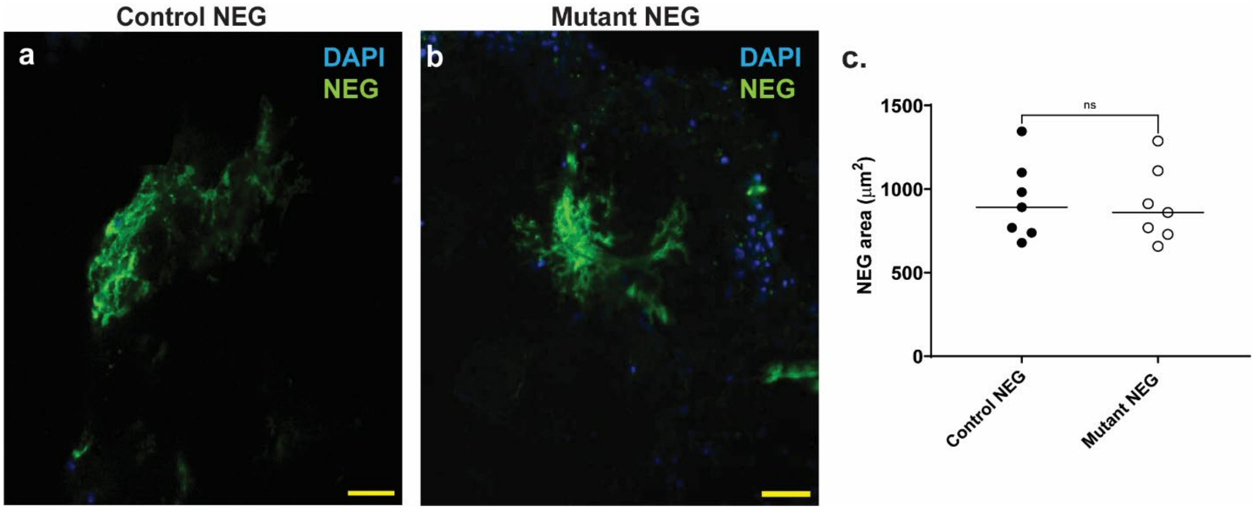


**Supplementary Figure S5:** NEGs show regular morphology in *drd* mutants. a. A single NEG in a 2-day old adult control brain. b. A single NEG in a 2-day old adult *drd* mutant brain. Nuclei were stained with DAPI (blue) and anti-FLAG was used to label NEGs (green). Scale bar=10 μm. c. Comparison of NEG cell areas in mutant and control. n= 7 cells from 9 control brains and 7 cells from 9 mutant brains, Mann-Whitney test, p=0.7.

**Supplementary Figure S6**

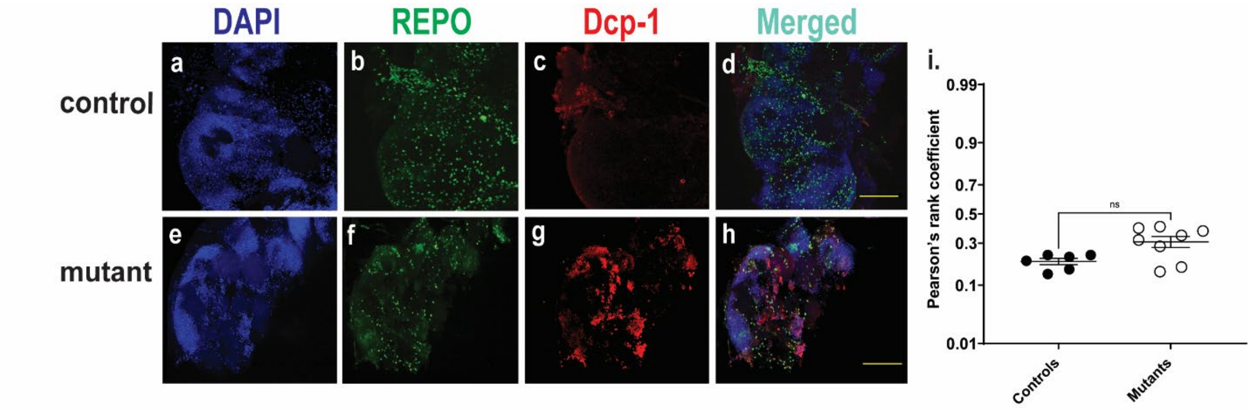


**Supplementary Figure S6:** Glial cells show less colocalization with apoptotic cells in 0-day old mutants. (a-h) localization of Dcp-1 with respect to REPO in brain cortex. Scale 100µm. (i) Pearson's rank coefficient for controls= 0.20, mutants= 0.31, n= 6 control and 8 mutant brains, Mann-Whitney test, p= 0.05.

**Supplementary Figure S7**

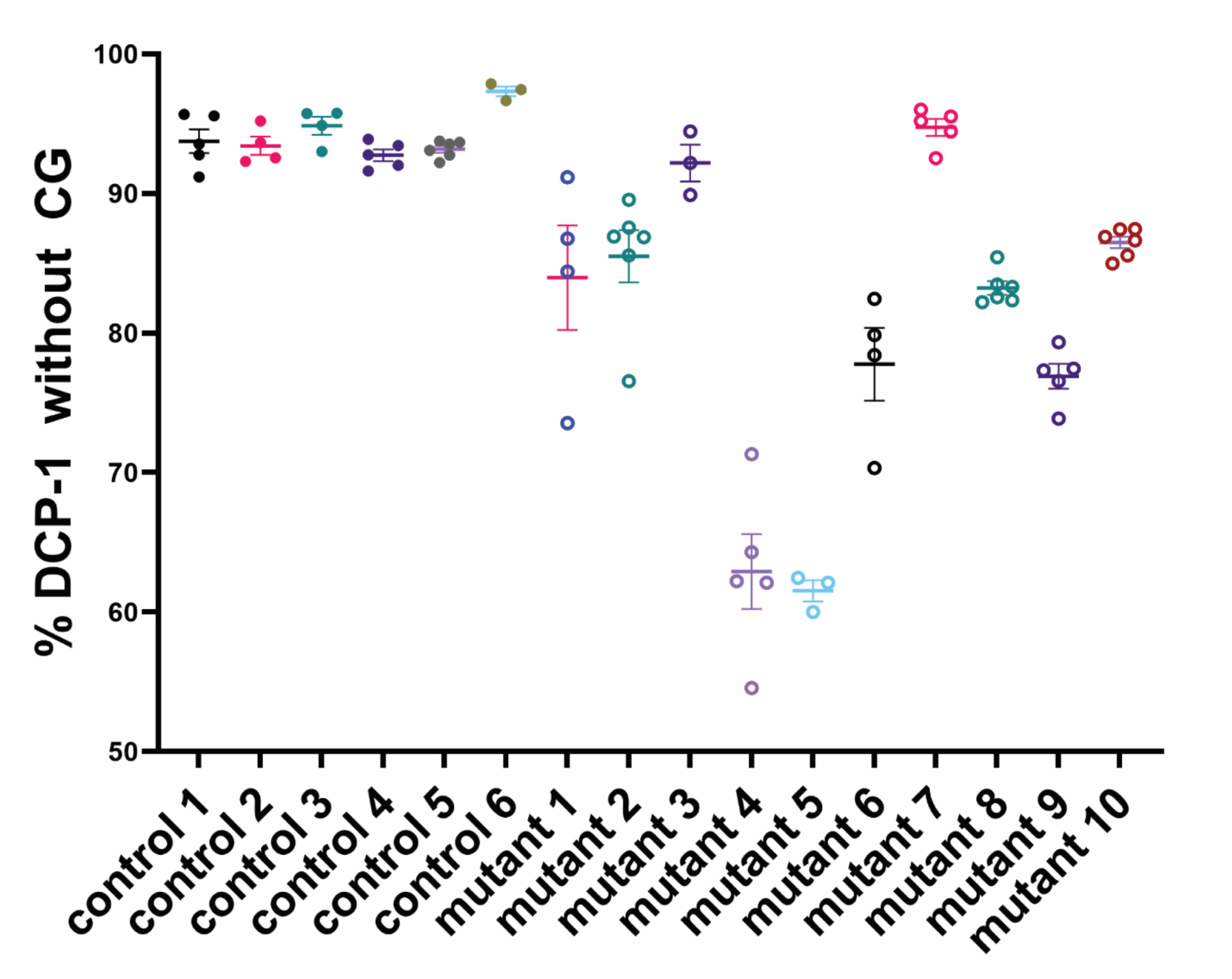


**Supplementary Figure S7:** The majority of Dcp-1 staining lies outside of the CG network. Object based colocalization tests show 56%-92% of Dcp-1 puncta outside of the CG network in mutants compared to 90% in controls. n= 6 controls and 10 mutant brains, Mann-Whitney test, p=0.55.

**Supplementary Table S1**

DAPI numbers and %area covered within CG

| Image | Area of overlap (and) | Area of no overlap (subtract) | No. of DAPI stained cells | Total area | %volume of DAPI with CG |
|---|---|---|---|---|---|
| **mutant 1** | 6647.53 | 4468.86 | 733 | 9454.34 | 71.52% |
| **mutant 2** | 6129.34 | 4546.77 | 740 | 9234.43 | 64.32% |
| **mutant 3** | 6997.54 | 4555.65 | 820 | 9134.43 | 73.56% |
| **mutant 4** | 6376.56 | 4467.89 | 732 | 8456.43 | 72.78% |
| **mutant 5** | 6678.54 | 4845.76 | 721 | 8767.65 | 75.32% |
| **mutant 6** | 6213.34 | 4623.55 | 721 | 9124.56 | 67.45% |
| **mutant 7** | 6287.54 | 4467.54 | 678 | 9345.65 | 67.78% |
| **mutant 8** | 6657.4 | 3594.76 | 545 | 9234.54 | 67.67% |
| **mutant 9** | 6367.45 | 3655.78 | 723 | 9123.54 | 61.32% |
| **mutant 10** | 7004.76 | 3556.32 | 756 | 10094.87 | 60.95% |
| control 1 | 8324.54 | 411.65 | 2923 | 9123.56 | 92.56% |
| control 2 | 9345.56 | 567.89 | 2745 | 9656.65 | 96.67% |
| control 3 | 9324.34 | 387.5 | 2756 | 9756.65 | 95.31% |
| control 4 | 9342.34 | 426.87 | 2967 | 9921.56 | 92.98% |
| control 5 | 9356.87 | 511.34 | 2776 | 9575.45 | 97.45% |
| control 6 | 9154.78 | 398.54 | 2667 | 9598.56 | 93.67% |

**Supplementary Table S1**. OBC analysis between DAPI and CG and counting of DAPI stained cells within CG network. Mean percentage of DAPI stained nuclei within the CG network is 95.42% for controls and 38.02% for mutants. The mean DAPI numbers within CG network for controls and mutants are 723 and 2774 respectively.

**Supplementary text S1: Macro code for volume analysis**

```
"Measure Stack" {
   run("Clear Results");   // First, clear the results table

   // loop through each slice in the stack. Start at n=1 (the first slice),
   // keep going while n <= nSlices (nSlices is the total number of slices in the stack)
   // and increment n by one after each loop (n++)
   for (n=1; n<=nSlices; n++) {
     setSlice(n);  // set the stack's current slice to n
     run("Measure");   // Run the "Measure" function in ImageJ
   }

   // Create a variable that we will use to store the area measured in each slice
   totalArea = 0;
   // Loop through each result from 0 (the first result on the table) to nResult (the total number of
results on the table)
   for (n=0; n < nResults; n++)
   {
     totalArea += getResult("Area",n);   // Add the area of the current result to the total
   }
   // Get the calibration information from ImageJ and store into width, height, depth, and unit
variables.
   // We will only be using depth and unit
   getVoxelSize(width, height, depth, unit);
   // Calculate the volume by multiplying the sum of area of each slice by the depth
   volume = totalArea*depth;
   // Print the result of the volume calculation to the log
   print(volume + " " + unit + "^3");
}
```
